# Mutant *Ppbbx24-del* gene positively regulates light-induced anthocyanin accumulation in the red pear

**DOI:** 10.1101/2023.05.19.541476

**Authors:** Shuran Li, Chunqing Ou, Fei Wang, Yanjie Zhang, Omayma Ismail, Yasser S.G. Abd Elaziz, Sherif Edris, Shuling Jiang, He Li

## Abstract

Anthocyanins are pigments and nutrients in red pears regulated by BBX family genes. Herein, we characterized a 14-nucleotide deletion mutation in the coding region of the *PpBBX24* gene from ‘Red Zaosu’ pear (*Pyrus pyrifolia* White Pear Group), named *Ppbbx24-del*. Genetic and biochemical approaches were used to compare the roles of PpBBX24 and Ppbbx24-del in anthocyanin accumulation. *Ppbbx24-del* played a positive role in anthocyanin biosynthesis of the ‘Red Zaosu’ pear peel by light treatment. Functional analyses based on overexpression in tobacco and transient overexpression in pear fruit peels showed that *Ppbbx24-del* promoted anthocyanin accumulation. Cyanidin and peonidin were major differentially expressed anthocyanins, and transcript levels of some structural genes in the anthocyanin biosynthesis pathway were significantly increased. Protein interaction assays showed that PpBBX24 was located in the nucleus and interacted with PpHY5, whereas Ppbbx24-del was colocalized in the nucleoplasm and did not interact with PpHY5. PpHY5 and Ppbbx24-del had positive regulatory effects on the expression of *PpCHS*, *PpCHI*, and *PpMYB10* when acting alone, but had cumulative effects on gene activation when acting simultaneously. Alone, PpBBX24 had no significant effect on the expression of *PpCHS*, *PpCHI*, or *PpMYB10*, whereas it inhibited the activation effects of PpHY5 on downstream genes when it existed with PpHY5. Our study demonstrated that mutant Ppbbx24-del positively regulates the anthocyanin accumulation in pear. The results of this study clarify the mechanism and enrich the regulatory network of anthocyanin biosynthesis, which lays a theoretical foundation for *Ppbbx24-del* use to create red pear cultivars.

## Introduction

Pear (*Pyrus* spp.) fruits are popular among consumers because of their crisp and juicy taste (Liu et al., 2021c). They are divided into Oriental and Occidental pears depending on the place of origin, and can be also divided into green-, yellow-, brown-, and red-skinned pear cultivars (Gamble et al., 2006). Red pear fruits have a huge market demand because of their unique appearance, quality, and nutritional value (Sun et al., 2014). However, red pear cultivars are rare, but more common in Occidental pears (Bao et al., 2008). For crisp oriental pears, the red peel color is difficult to obtain and cultivars are rare (Bai et al., 2019a). Thus, the breeding of red pear cultivars is an important goal in China and worldwide (Sun et al., 2021a; Zhai et al., 2021).

Accumulation of anthocyanins in pear peels is the main determinant of red coloration in the fruit (Sun et al., 2014). Anthocyanin biosynthesis belongs to a branch of the flavonoid biosynthetic pathway (Fang et al., 2019a), particularly, the phenylpropionic acid and flavonoid synthesis pathways (Pervaiz et al., 2017). Two types of genes are involved in anthocyanin synthesis: structural and regulatory genes (Ye et al., 2017; Sun et al. 2021b; Alabd et al. 2022). Structural genes mainly encode for early enzymes in the anthocyanin biosynthetic pathway, such as phenylalanine ammonia-lyase (PAL), chalcone synthase (CHS), chalcone isomerase (CHI), flavanone3-hydroxylase (F3H), and late-pathway enzymes including dihydroflavonol 4-reductase (DFR), anthocyanidin synthase (ANS), and flavonoid 3-O-glycosyl-transferase (UFGT) (Han et al. 2020; Yin et al. 2021). Their expression levels are positively correlated with anthocyanin content in the peach (Ye et al. 2017b), apple (Fang et al. 2019a; Hu et al. 2021), pear (Ni et al. 2020; Alabd et al. 2022), grape (Su et al. 2023), and strawberry (Martinez-Rivas et al. 2023). On the other hand, regulatory genes are transcription factors that regulate anthocyanin biosynthesis pathways at the transcriptional level (Zhang et al. 2020). *MYB*, *bHLH*, and *WD40* were first identified as important regulatory genes involved in the synthesis of anthocyanins by forming the MBW (MYB-bHLH-WD40) complex (Chen et al. 2021; Zhang et al. 2023). Among the MBW complexes, MYB is one of the largest families of transcription factors, and several of its members can activate the expression of structural genes. Multiple *MYB* transcription factors including *MdMYBA* (Mercan et al. 2021), *MdMYB1* (Hu et al. 2021), *MdMYB2* (Jiang et al. 2022), *MdMYB10* (Wang et al. 2020), *MdMYB16* (Mao et al. 2021), and *MdMYB110a* (Sato et al. 2019) have been identified in apples and are related to anthocyanin synthesis. *MdMYB10* is one of the most critical transcription factors involved in regulating anthocyanin synthesis in fruit (Wang et al., 2023). *MYB10* has also been found to be widely present in Rosaceae fruits, including peach (Karppinen et al., 2021), cherry (Calle et al., 2021), pear (Wang et al., 2020), strawberry (Castillejo et al., 2020). Numerous reports have identified many regulatory genes, including *MdBT2* (An et al., 2019; Ren et al., 2021), *VcNAC072* (Yang et al., 2019), *LcNAC13* (Jiang et al., 2019), *MdERF38* (An et al., 2020b), *FvMAPK3* (Mao et al., 2022), *MdMPK6* (Xing et al., 2023) and *PpERF9* (Ni et al., 2023).

Recently, several *MYB* transcription factors associated with anthocyanin synthesis have been identified in pears (Yao et al., 2017; Yao et al., 2023). The methylation level of the pear *MYB10* promoter can affect the expression of this gene, which may be related to the red coloration of the fruit (Qian et al., 2014). The expression of the *MYB10* gene in ‘Hongsha’ (Feng et al., 2015) and ‘Yunhongli No.1’ pears (Wang et al., 2020) was promoted by light, regulating the coloration of the peel effectively. Additionally, the PyMYB114/PyMYB10 complex promotes anthocyanin biosynthesis in the pear peel (Liu et al., 2021a). PpMYB140 can directly inhibit the expression of structural genes and suppress anthocyanin accumulation by competing with PpMYB114 for binding to bHLH3 (Ni et al., 2021). Numerous studies have shown that many *MYB* transcription factors play a regulatory role in the biosynthesis of anthocyanin, such as *PpMYB12b* (Zhai et al., 2019), *PpMYB17* (Premathilake et al., 2020), *PbMYB26* (Zhao et al., 2021a), *PyMYB73* (Yao et al., 2023) and *PbMYB120* (Song et al., 2020). Furthermore, many transcription factors affect anthocyanin biosynthesis in pears. PpHY5 directly binds to the G-box of the *PyMYB10* promoter to induce its expression, thereby promoting anthocyanin accumulation (Tao et al., 2018; Bai et al., 2019b; Wang et al., 2020). *PpbHLH64* (Tao et al., 2020), *PbWRKY75* (Zhai et al., 2021), *Pp4ERF24* and *Pp12ERF96* (Ni et al., 2019) positively regulate anthocyanin biosynthesis. The biological function of these genes is mainly caused by the difference in expression between red and green fruits, rather than by the difference in the sequences of the gene coding and promoter regions.

BBX (B-box) proteins are zinc finger structural proteins involved in a variety of photoregulated plant physiological processes, including photomorphogenesis, circadian rhythms, coloration, shade avoidance, and flowering (Job et al., 2018; Brusch et al., 2020; Yadav et al., 2020). There are 32, 30, and 64 *BBX* family members identified in *Arabidopsis*, rice, and apple, respectively (Huang et al., 2012; Liu et al., 2017; Lyu et al., 2020a). *BBX24* is a member of the *BBX* family, also known as the salt tolerance gene (*STO*), which was first identified in *Arabidopsis* and is mainly involved in UV-B signal transduction (Lippuner et al., 1996; Lyu et al., 2020b). In apples, BBX24 inhibits anthocyanin synthesis by interacting with HY5 (Fang et al., 2019a). BBX24 also regulates corolla coloring in petunias by interacting with HY5 (Fu et al., 2020). *GhDR* is homologous to *AtBBX24* and is associated with the accumulation of anthocyanins in cotton (Wang et al., 2022). There are 37 members of the *BBX* family that have been identified in pears (Talar et al., 2021). Although there have been reports that BBX16, BBX18, and BBX21 interact with HY5 and affect the biosynthesis of pear anthocyanins, there are few reports on the function of *PpBBX24* (Bai et al., 2019a; Bai et al., 2019b).

’Red Zaosu’ is a pear cultivar that originated from the discovery of a red mutant of ‘Zaosu’ (Li et al., 2021b). The mutant inherits better fruit quality and strong stress resistance of ‘Zaosu’ and is also the only type of crisp fruit among many red mutants in production (Tao et al., 2018). In the previous study, we identified a novel 14 nucleotide deletion mutation from the red pear mutant ‘Red Zaosu;’ the deletion variant is located at the coding region of the *PpBBX24* gene, which causes a coding frameshift and early termination (Ou et al., 2020). This deletion mutation is named *Ppbbx24-del*, and it is the only mutation associated with red variation identified in the mutant. Therefore, to study the molecular mechanism of its regulation on the coloration of ‘Red Zaosu,’ in this work, the expression and function of the *Ppbbx24-del* gene were analyzed for the first time by applying various approaches. Our work resulted in a regulatory model for Ppbbx24-del and PpBBX24 proteins in the anthocyanin accumulation process, which enriches the regulatory network of plant anthocyanin biosynthesis and provides theoretical support for creating a new red pear cultivar using gene editing technology.

## Results

### Fruit anthocyanin levels and expression analysis of *PpBBX24* and *Ppbbx24-del* under light conditions

The bags were removed 21 days before ‘Zaosu’ and ‘Red Zaosu’ fruit ripening. The color of the ‘Red Zaosu’ pear peel became darker gradually with the extension of illumination. After 21 days of natural light, the peel of ‘Red Zaosu’ was the most stained and the anthocyanin content was 56 times that of the ‘Zaosu’ pear peel, whereas the ‘Zaosu’ fruit barely accumulated anthocyanins throughout the treatment (Figure 1A and B).

**Figure 1.**
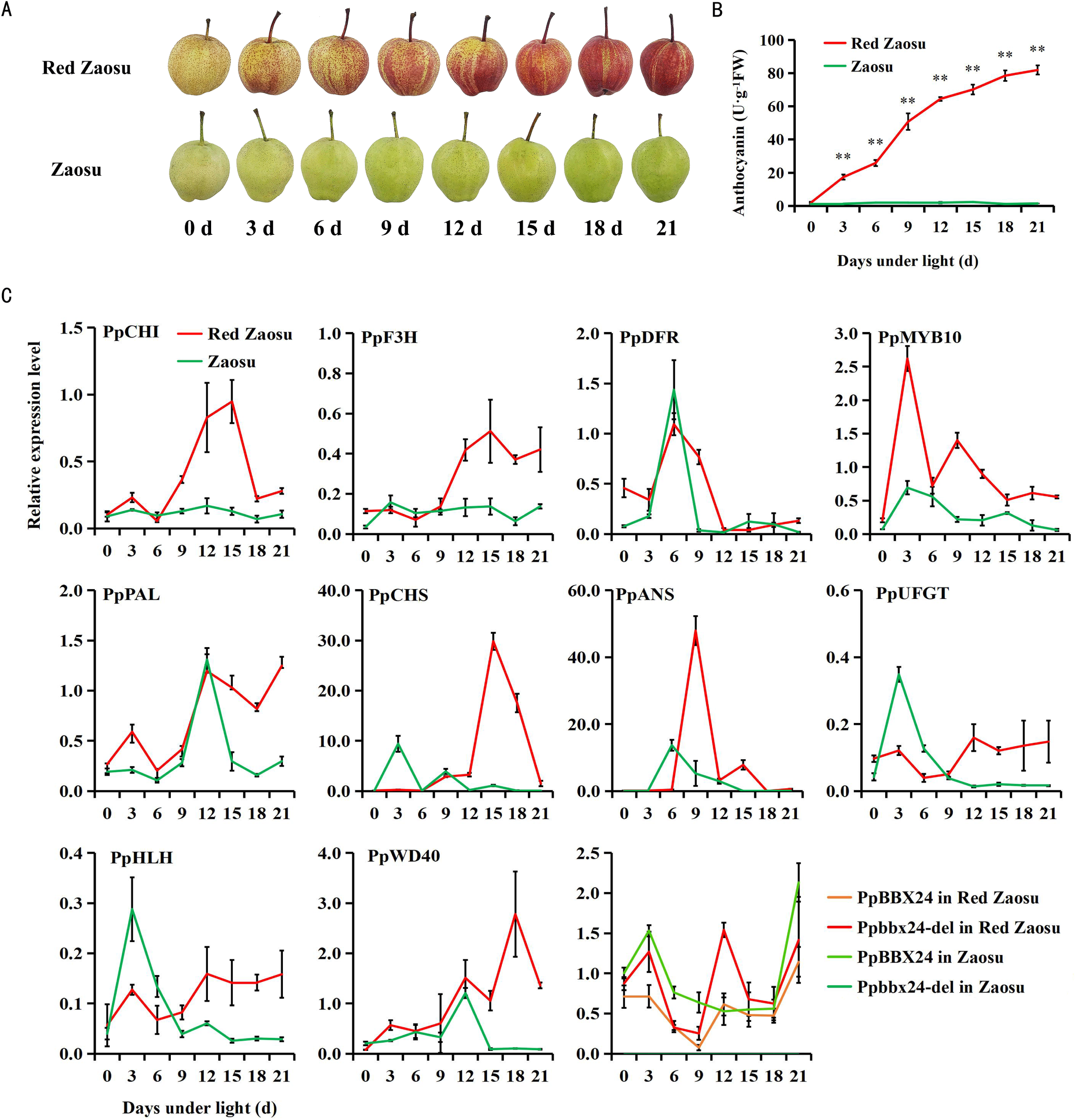
Fruit anthocyanin levels and expression analysis of *PpBBX24* and *Ppbbx24-del* under light. A, Effects of light on appearance of ‘Zaosu’ and ‘Red Zaosu’ pears. B, Analysis of anthocyanin contents at different illumination times. C, Transcript levels of structural genes in the anthocyanin biosynthesis pathway and *PpBBX24* and *Ppbbx24-del* after light treatment. Values are represented as the mean ± standard deviation of three independent biological replicates. Asterisks indicate significant differences (ip≤0.05, iip≤0.01), the same below.

To clarify the expression difference of *PpBBX24* and *Ppbbx24-del* genes in ‘Zaosu’ and ‘Red Zaosu’ pear peels, their relative expression levels were detected by RT-qPCR in the two varieties after bag-picking. In a previous study, we found that *Ppbbx24-del* existed in ‘Red Zaosu’ and its red progeny, showing a heterozygous state (Ou et al., 2020). However, ‘Zaosu’ and its green progeny did not have this deletion mutation, indicating that the *PpBBX24* gene in ‘Red Zaosu’ comprised both deletion (*Ppbbx24-del*) and normal (*PpBBX24*) types, while there was only normal (*PpBBX24*) type in ‘Zaosu.’ Compared with *PpBBX24*, *Ppbbx24-del* lost 14 bp, and the first six bases of the deletion site were completely consistent with the six bases after the mutation site (Figure S1). The primers designed across the deletion region were not sufficiently specific to distinguish the *Ppbbx24-del* gene. However, the primers designed for the mutated region could detect the expression of the normal-type *PpBBX24* gene (Table S1). Thus, we detected the co-expression levels of *PpBBX24* and *Ppbbx24-del* genes in ‘Red Zaosu’ by using primers designed across the mutation region, then subtracted the expression of *PpBBX24* with these values to obtain the expression levels of the mutant gene *Ppbbx24-del*. The results showed that the effect of light treatment on the expression of the *PpBBX24* gene was not significant in the peel of the ‘Zaosu’ pear. Although the expression of *PpBBX24* and *Ppbbx24-del* genes fluctuated slightly, there was no obvious regularity in the ‘Red Zaosu’ pear peel (Figure 1C).

Studies have shown that *PpCHI*, *PpF3H*, *PpDFR*, *PpMYB10*, *PpPAL*, *PpCHS*, *PpANS*, *PpUFGT*, *PpHLH*, and *PpWD40* are key structural genes in the anthocyanin biosynthesis pathway (An et al., 2021). They play important roles in the synthesis of anthocyanins in plants. To further clarify the expression difference of the above genes in the peel of the ‘Zaosu’ and ‘Red Zaosu’ pear, their relative expression levels were detected by RT-qPCR in the two cultivars after bag removal. After light treatment, the transcript levels of most anthocyanin structural genes in the ‘Red Zaosu’ pear peel were significantly higher than that in the ‘Zaosu’ pear peel (Figure 1C). These results indicate that *Ppbbx24-del* responds to light and may be related to anthocyanin accumulation.

### The subcellular localization of PpBBX24 and Ppbbx24-del in the pear

Preliminary results showed that due to early encoding termination, Ppbbx24-del lacks a nuclear localization sequence (NLS) compared with PpBBX24 (Ou et al., 2020). Therefore, the subcellular localizations of PpBBX24 and Ppbbx24-del may differ. In this study, 35S:PpBBX24-EGFP and 35S:Ppbbx24-del-EGFP recombinant vectors were constructed. After transient infection of the onion epidermis, the PpBBX24 fluorescent protein signal mainly appeared in the nucleus, whereas the Ppbbx24-del fluorescent protein signal appeared in both the nucleus and cytoplasm. In contrast, EGFP alone was found throughout the cell, indicating that the subcellular localization of Ppbbx24-del was different from that of PpBBX24 (Figure 2).

**Figure 2.**
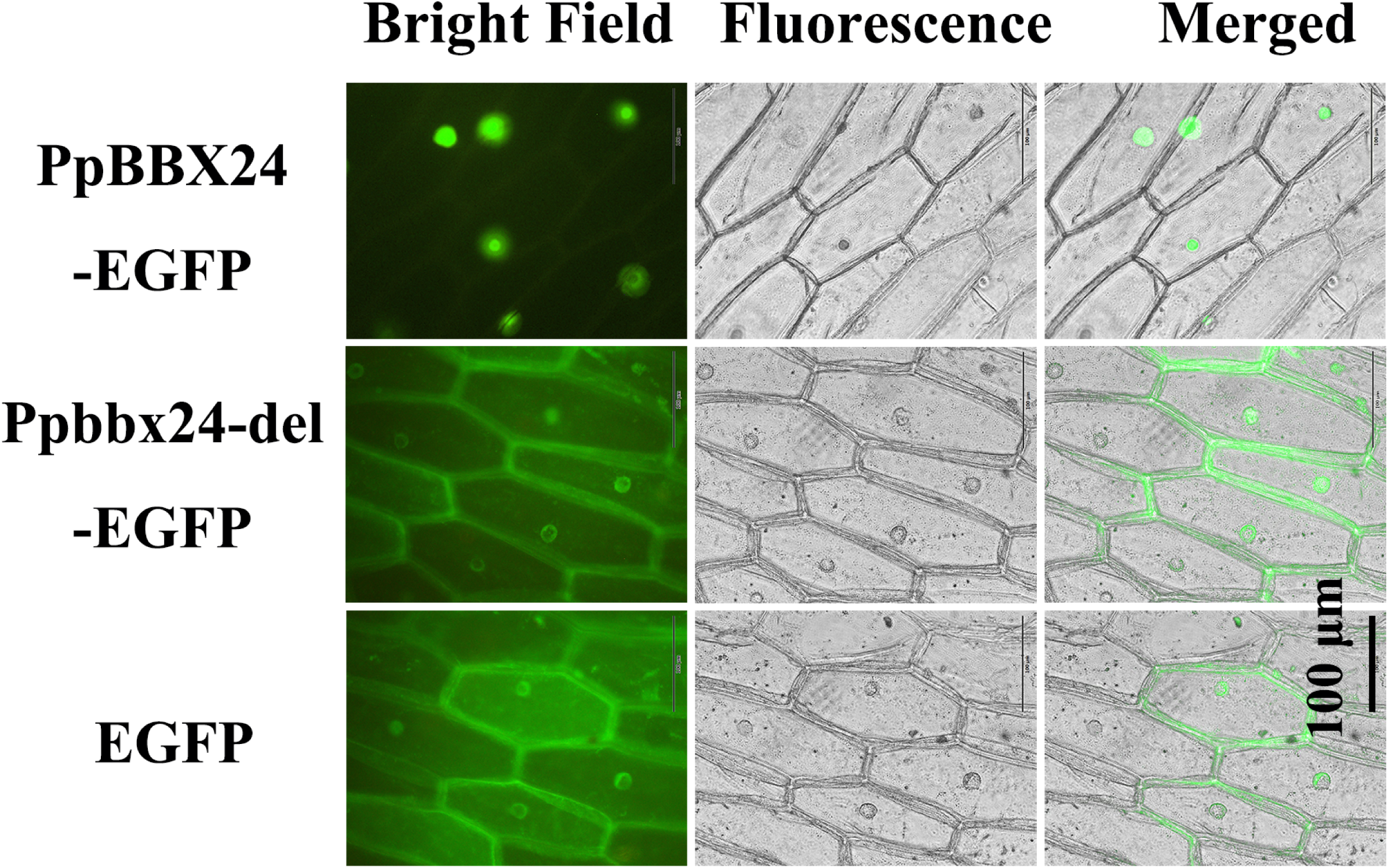
Nuclear localization of PpBBX24 and Ppbbx24-del in onion epidermis cells.

### *Ppbbx24-del* promotes anthocyanin accumulation in tobacco and pear

To elucidate the functions of *PpBBX24* and *Ppbbx24-del*, overexpression vectors of the two genes were constructed and transformed into tobacco plants using *Agrobacterium*. All transgenic lines were identified and tested using PCR (primers are shown in Table S1). Target bands of approximately 706 and 720 bp were amplified in *Ppbbx24-del* and *PpBBX24* transgenic tobacco, respectively. The target bands were amplified using a bacterial solution containing an overexpression vector. No target bands were detected in the wild-type tobacco. The results showed that transgenic plants overexpressing *Ppbbx24-del* and *PpBBX24* were generated (Figure 3A). To further clarify the effects of *Ppbbx24-del* on the coloration of tobacco flowers, we examined the phenotypic changes in the transgenic plants in detail. Most *Ppbbx24-del* transgenic lines displayed an anthocyanin-rich phenotype characterized by visible accumulation of these pigments in flowers. The presence of anthocyanins was particularly evident in the corolla, which displayed a deep red coloration with the colored area extending inside the corolla tube (Figure 3B). Morphology and size were monitored in flowers from *PpBBX24* transgenic tobacco and compared with those of wild-type flowers, and no significant differences were observed. Biochemical analysis confirmed that more than 2.2 times anthocyanin levels were present in the flowers of *Ppbbx24-del* transgenic plants than in *PpBBX24* transgenic plants and wild-type tobacco (Figure 3C). These results indicate that *Ppbbx24-del* can promote the coloring of tobacco flowers and participate in the accumulation of anthocyanin under normal light conditions, while the *PpBBX24* gene cannot promote the accumulation of anthocyanin in tobacco flowers.

**Figure 3.**
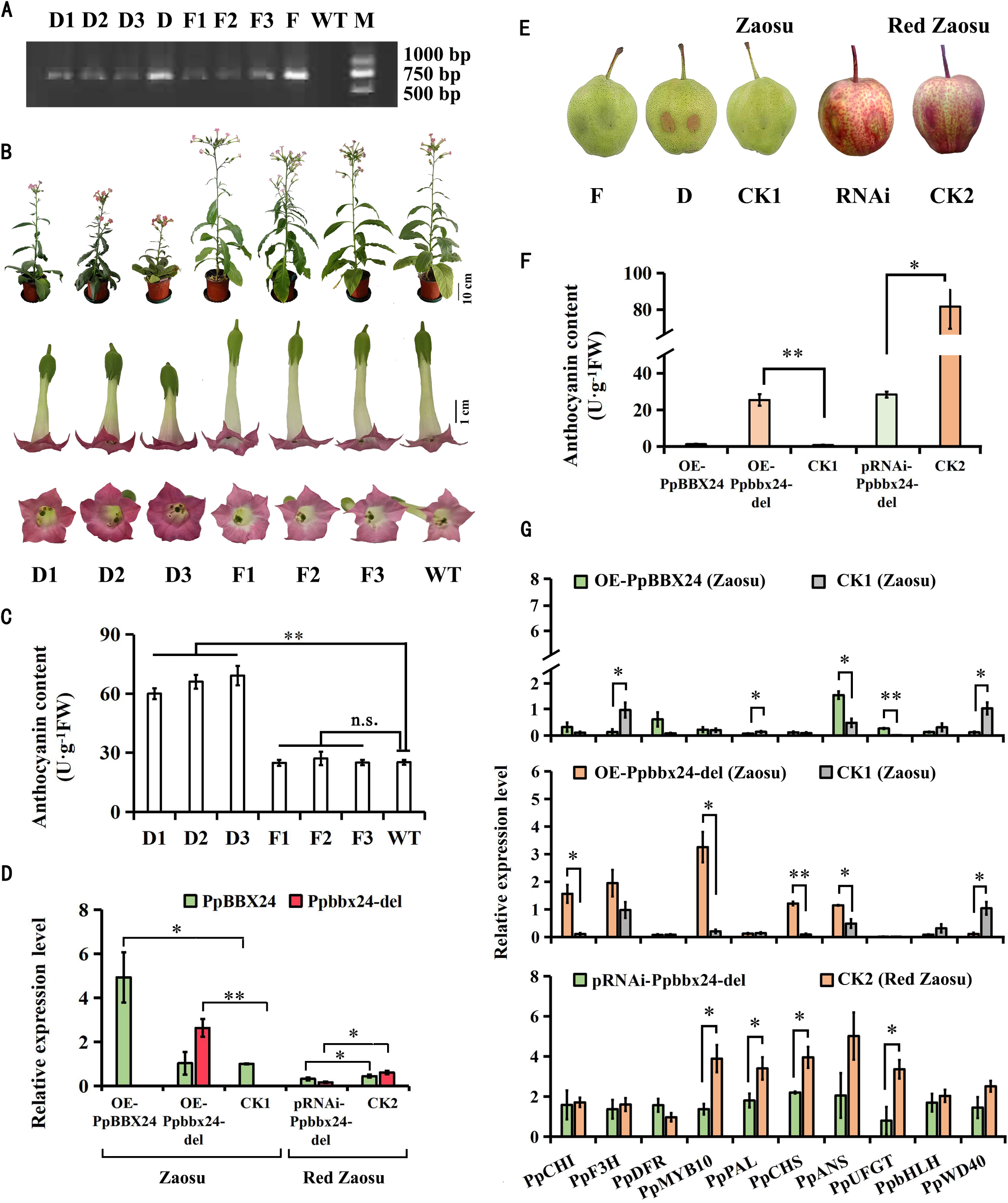
Effects of *PpBBX24* and *Ppbbx24-del* genes on anthocyanin accumulation in tobacco plants and pear peels. A, Presence of transgene in transgenic tobacco lines confirmed by PCR amplification; D1–D3 were OE-*Ppbbx24-del* lines; D was a control for *Ppbbx24-del* gene; F1–F3 were OE-*PpBBX24* lines; F was a control for *PpBBX24* gene; WT was a wild-type tobacco; and M was a 2000 bp DNA marker. B, Three independent *Ppbbx24-del*-overexpression transgenic lines, three independent *PpBBX24*-overexpression transgenic lines and wild-type tobacco plants and flowers in natural growth states. C, Anthocyanin contents in D1–D3, F1–F3, and WT tobacco flowers. D, Relative *PpBBX24* and *Ppbbx24-del* transcript levels in fruit transiently overexpressing *PpBBX24*, *Ppbbx24-del*, and RNAi. E, Pear fruit anthocyanin accumulation in the transient overexpression of *PpBBX24*, *Ppbbx24-del*, and RNAi after light treatment; F was transiently overexpressing *PpBBX24*; D was transiently overexpressing *Ppbbx24-del*; CK1 was the empty control of ‘Zaosu’ pear; RNAi was ‘Red Zaosu’ pear fruit in which *Ppbbx24-del* was transiently interference; and CK2 was empty control of ‘Red Zaosu’ pear. F, Anthocyanin contents around infiltrated sites of fruit peel transiently overexpressing *PpBBX24*, *Ppbbx24-del*, and RNAi. G, Transcript levels of *PpMYB10* and structural genes in the anthocyanin biosynthesis pathway in pear peel transiently overexpressing *PpBBX24*, *Ppbbx24-del*, and RNAi. n.s. indicates no significant difference.

Transient expression of target genes is a simple and effective method to analyze biological function and has been widely used in the study of gene function in plants such as apples, peaches, and pears (Dang et al., 2021; Fan et al., 2023; Ni et al., 2023). To further clarify the function of *PpBBX24* and *Ppbbx24-del* genes on the coloration of pear peel, *A. tumefaciens* GV3101 with the pRI101-*PpBBX24*, pRI101-*Ppbbx24-del*, and RNAi-*Ppbbx24-del* vectors were injected into ‘Zaosu’ and ‘Red Zaosu’ pear fruit. The results showed after overexpression, the expression of *PpBBX24* and *Ppbbx24-del* genes in the ‘Zaosu’ pear peel were significantly higher than those in the control (CK1). After interference, the expression of *Ppbbx24-del* genes in the ‘Red Zaosu’ pear peel were significantly lower than those in the control (CK2) (Figure 3D). After light treatment, the OE-*Ppbbx24-del* (D) pear peels appeared red, whereas the OE-*PpBBX24* (F) and CK1 pear peels were green. After *Ppbbx24-del* interference (RNAi), the pear peels turned green (Figure 3E). Spectrophotometric analyses confirmed that OE-*Ppbbx24-del* promoted anthocyanin accumulation; the anthocyanin content was about 33.7 times that of CK1. However, there was no significant difference in the anthocyanin content between OE-*PpBBX24* and CK1. RNAi reduced anthocyanin accumulation compared to that in CK2 (Figure 3F).

To further clarify the differences in the expression of anthocyanin structural genes after injection, transcript levels at the injection site were determined using RT-qPCR. These analyses revealed that *PpCHI*, *PpMYB10*, *PpCHS*, and *PpANS* were upregulated to various degrees in OE-*Ppbbx24-del*, and the expression levels of *PpMYB10*, *PpCHS*, and *PpCHI* were 10 times higher than those in CK1. *PpMYB10*, *PpPAL*, *PpCHS*, and *PpUFGT* were downregulated to varying degrees in RNAi-*Ppbbx24-del* (Figure 3G). These results indicate that *PpbbX24-del* promoted anthocyanin accumulation in ‘Red Zaosu’ pear peels under light conditions, whereas *PpBBX24* had no significant effect on anthocyanin accumulation.

### Anthocyanin content and transcript levels of structural genes in the anthocyanin biosynthesis pathway were different in green and red pear peels

To further analyze the mechanism of *Ppbbx24-del* in the coloration of the ‘Red Zaosu’ pear peel, transcriptomic sequencing and metabolome analysis were performed on the young leaves and fruit peels of ‘Zaosu,’ ‘Huangguan,’ and ‘Red Zaosu’ pears, as well as the red and green pools constructed by the young leaves of ‘Red Zaosu × Huangguan’ F1 plants. The anthocyanin biosynthesis pathway in pears was constructed by comparing the expression of structural genes and changes in metabolite components in the anthocyanin biosynthesis pathway in eight samples. Based on the metabolomic analysis results, Peonidin-3-O-arabinoside, Peonidin-3-O-rutinoside, Cyanidin-3,5-O-diglucoside, and Cyanidin-3-O-(6’’-O-malonyl) glucoside were the main metabolites in the pear peel and determined the fruit coloration. The green sample contained almost no anthocyanins (Figure 4).

**Figure 4.**
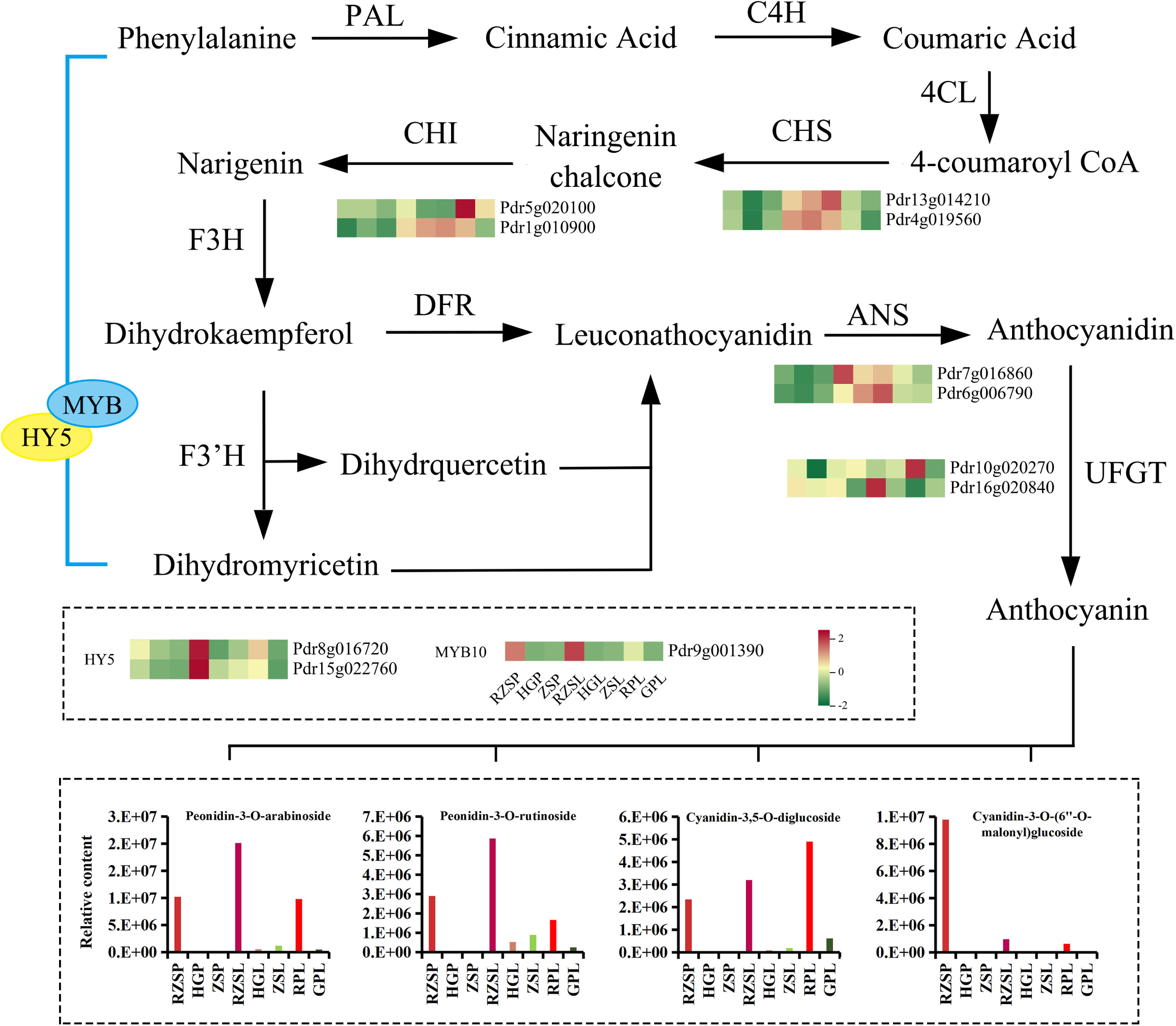
Anthocyanin biosynthesis pathway in pears. The column charts are drawn according to the FPKM values from the transcriptome datasets. Columns and rows in the column charts represent samples and genes, respectively. The heatmaps are drawn according to the results from the metabolome datasets. Columns and rows in the heatmaps represent samples and metabolites, respectively.

Through light treatment and transient overexpression, we found that the *Ppbbx24-del* gene promoted the coloration of ‘Red Zaosu’ pear peels, and the transcript levels of some structural genes also increased significantly in red samples. To further compare the expression of these genes in samples of different colors, we compared the transcriptome sequencing results. The expression of *PpCHS* (Pdr13g014210, Pdr4g019560), *PpCHI* (Pdr5g020100, Pdr5g020100), *PpANS* (Pdr7g016860, pdrgg6006790), and *PpUFGT* (Pdr10g020270, Pdr16g020840) in the red samples may be regulated by Ppbbx24-del. In addition, the *MYB* and *HY5* transcription factors play an essential role in regulating anthocyanin biosynthesis (Job et al., 2018; Liu et al., 2021a; Sun et al., 2023); therefore, we also analyzed the transcript levels of these genes. The expression of *PpMYB10* (Pdr9g001390) and *PpHY5* (Pdr8g016720 and Pdr15g022760) in the red samples increased significantly and may be regulated by Ppbbx24-del.

### PpBBX24 and Ppbbx24-del have different interaction modes with PpHY5

Studies on plants have shown that many BBX proteins affect anthocyanin accumulation through interaction with HY5 such as in *Arabidopsis*, pears, and apples (Job et al., 2018; An et al., 2020; Fu et al., 2020; Liu et al., 2021b; Liu et al., 2022). We also found that *PpHY5* was highly expressed in red samples in the transcriptome sequencing results. Therefore, we used BiFC, yeast two-hybrid, and pull-down assays to examine whether PpBBX24 and Ppbbx24-del physically interacted with PpHY5. In the BiFC assays, yellow fluorescence was observed when PpBBX24 was co-expressed with PpHY5, whereas the co-expression of Ppbbx24-del and PpHY5 was not observed (Figure 5A), indicating that PpBBX24 interacted with PpHY5, but Ppbbx24-del did not. In yeast two-hybrid assays, only when the PpBBX24-BD (the full-length PpBBX24 sequence fused with pGBKT7 vector) was cotransformed with PpHY5-AD (PpHY5 fused with pGADT7 vector), there was growth on SD/-Trp/-Leu/-His/AbA/X-α-Gal medium and the colonies appear blue (Figure 5B). This also indicated that PpBBX24 interacted with PpHY5, but Ppbbx24-del did not. In the pull-down assays, only PpBBX24-His was pulled down by PpHY5-GST with a His antibody (Figure 5C), and only PpBBX24 interacted with PpHY5. Therefore, we believe that PpBBX24 can interact with PpHY5, whereas Ppbbx24-del cannot and that there may be functional differences between the two.

**Figure 5.**
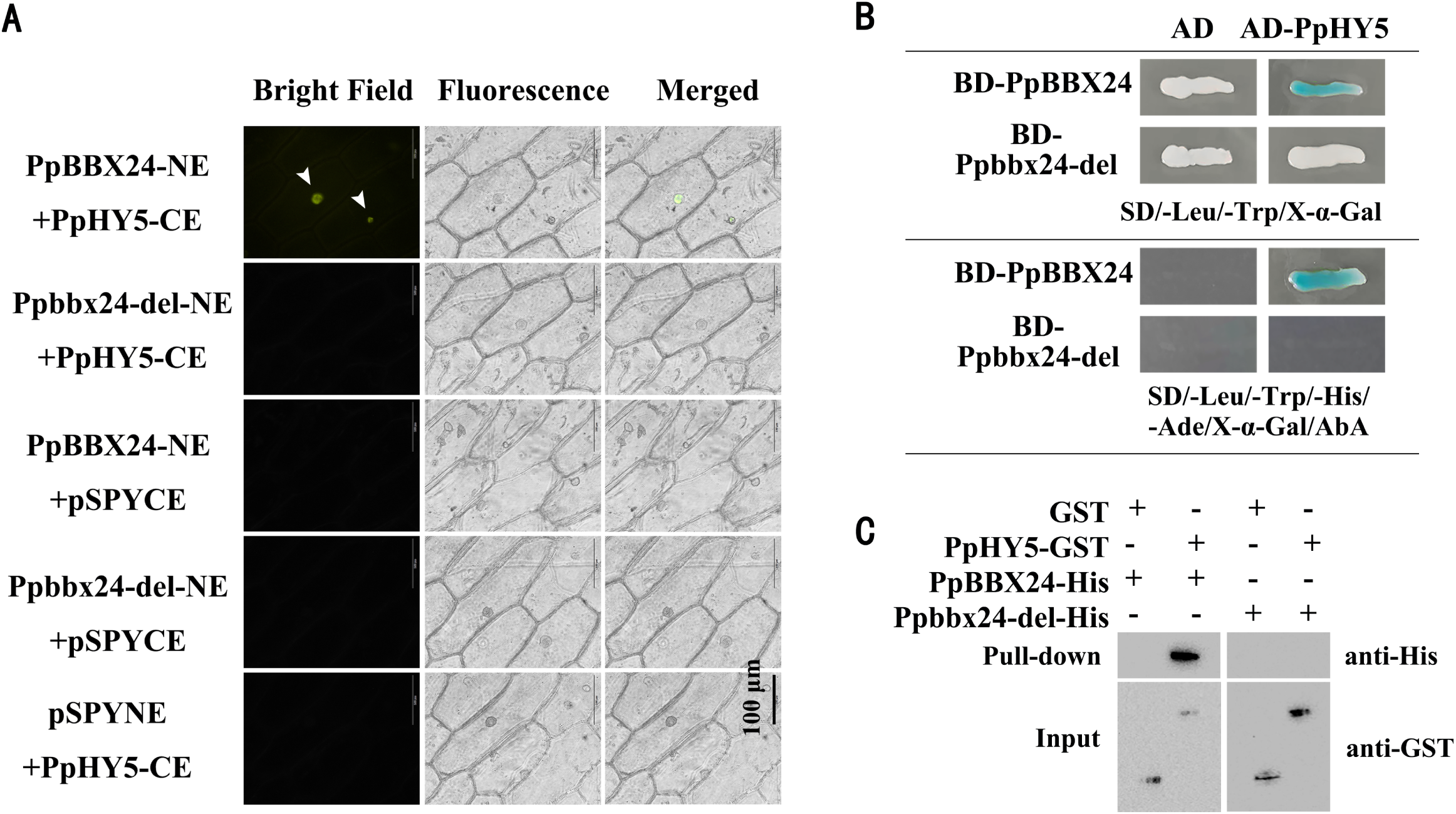
Physical interaction between PpBBX24, Ppbbx24-del, and PpHY5. A, Interaction between PpBBX24, Ppbbx24-del, and PpHY5 in vivo in the bimolecular fluorescence complementation (BiFC) assay. B, Interaction between PpBBX24, Ppbbx24-del, and PpHY5 in yeast two-hybrid assays. C, Pull-down assay performed by co-purifying recombinant PpBBX24-His (left) and Ppbbx24-del-His (right) fusion proteins with PpHY5-GST (column 1) and a GST empty vector (column 2). Western blotting with His antibody showed a pull-down of PpBBX24 and Ppbbx24-del by PpHY5-GST.

### PpBBX24 and Ppbbx24-del functions synergistically with PpHY5 to modulate *PpCHS*, *PpCHI*, and *PpMYB10* genes

In addition to *PpHY5*, we found that *PpCHS*, *PpCHI*, *PpANS*, *PpUFGT*, and *PpMYB10* were highly expressed in red samples in the transcriptome sequencing results, which may be involved in regulating the coloration of ‘Red Zaosu’ pear peels. The G-box domain has been reported to bind to BBX proteins (Gangappa et al., 2014; Fang et al., 2019a). To characterize the expression of these structural genes, we analyzed the promoter regions up to 1500–2000 bp upstream of the translation start site using the PlantCARE database. There was no difference in the promoter sequences of these genes between ‘Zaosu’ and ‘Red Zaosu,’, and there were multiple promoter cis-acting elements in both varieties, including the CAAT-box and TATA-box. Except for PpUFGT, the other five gene promoters contained 2–5 G-box motifs (Figure 6A).

**Figure 6.**
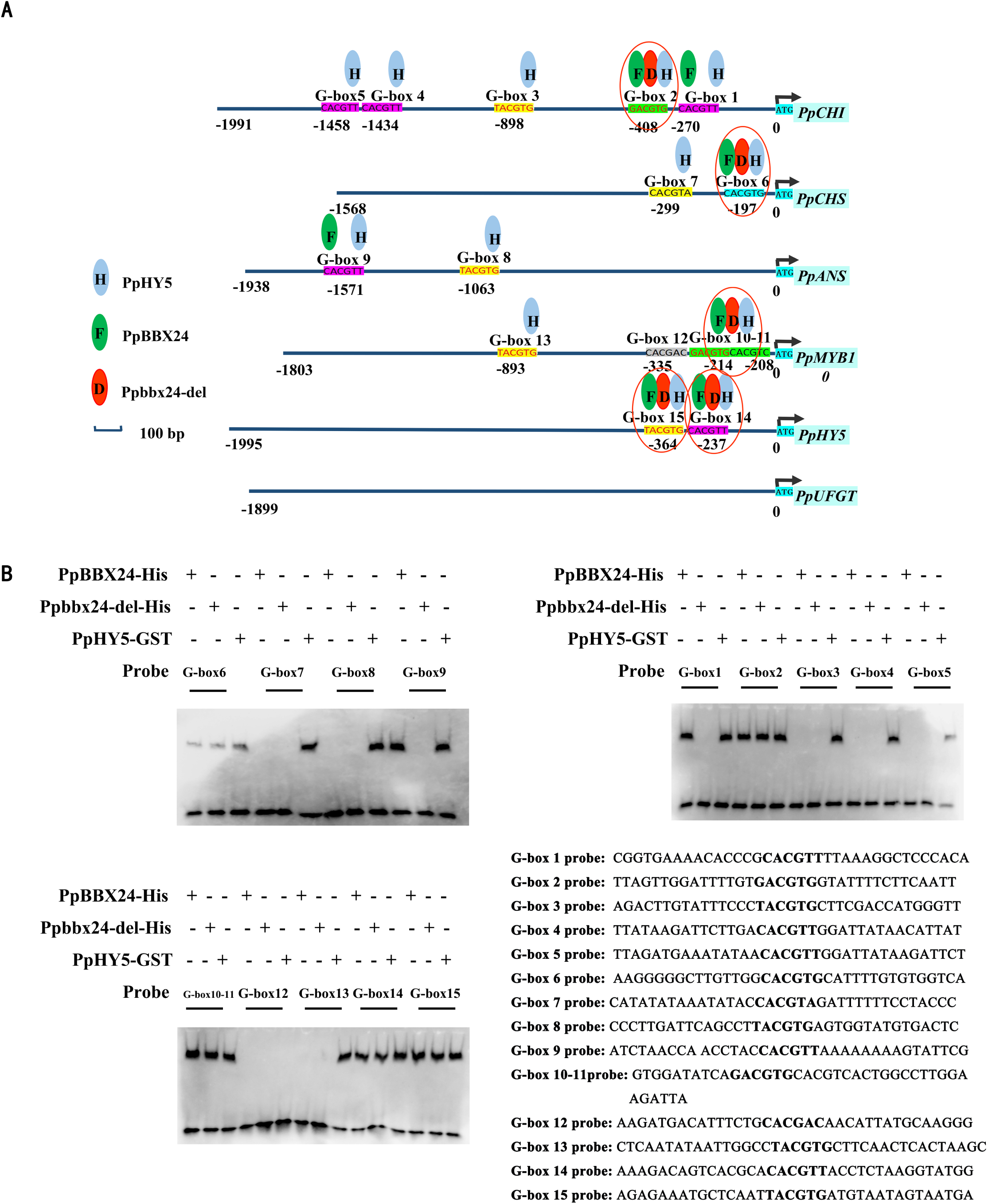
G-box motifs analysis of promoters of *PpCHI*, *PpCHS*, *PpANS*, *PpMYB10*, *PpHY5*, and *PpUFGT*. A, Results of G-box motifs in the promoter of *PpCHI*, *PpCHS*, *PpANS*, *PpMYB10*, *PpHY5*, and *PpUFGT*. B, Interactions of PpBBX24, Ppbbx24-del, and PpHY5 with G-box probes in EMSA test. C, G-box probes in promoters of *PpCHI*, *PpCHS*, *PpANS*, *PpMYB10*, and *PpHY5*.

Previous studies have shown that BBX24 and HY5 can bind to G-box motifs in the promoters of anthocyanin structural genes (Fang et al., 2019a; Xing et al., 2023), and analyses of promoter sequences have revealed G-box motifs in the promoters of *PpCHI*, *PpCHS*, *PpANS*, *PpHY5*, and *PpMYB10*. An EMSA assay was used to analyze the interactions. In the EMSA, PpBBX24, Ppbbx24-del, and PpHY5 all bound to the promoters of *PpCHI*, *PpCHS*, *PpHY5*, and *PpMYB10* (Figure 6B). In the light treatment, transient overexpression, and transcriptome sequencing assays, the transcript levels of some structural genes increased significantly in the red samples (Figure 1D, 3F, 4). To investigate whether Ppbbx24-del promotes anthocyanin production by directly regulating these genes, three genes with the most varied expression levels, *PpCHS*, *PpCHI,* and *PpMYB10*, were selected. We conducted a rigorous EMSA to further verify the interactions between PpBBX24, Ppbbxx24-del, and the promoters of these genes. These results showed that PpBBX24 and Ppbbx24-del proteins could bind to the G-box motifs of the *PpCHS*, *PpCHI*, and *PpMYB10* promoters in vitro (Figure 7B). The results of the yeast one-hybrid assays were consistent with those of the EMSA (Figure 7B). Because HY5 can transmit light signals to downstream components (Fang et al., 2019a), we confirmed that PpHY5 could also bind to the G-box motifs of the *PpCHS*, *PpCHI*, and *PpMYB10* promoters using EMSA and yeast one-hybrid assays. These experiments indicate that PpBBX24, Ppbbx24-del, and PpHY5 directly bind to the promoters of *PpCHS*, *PpCHI*, and *PpMYB10*.

**Figure 7.**
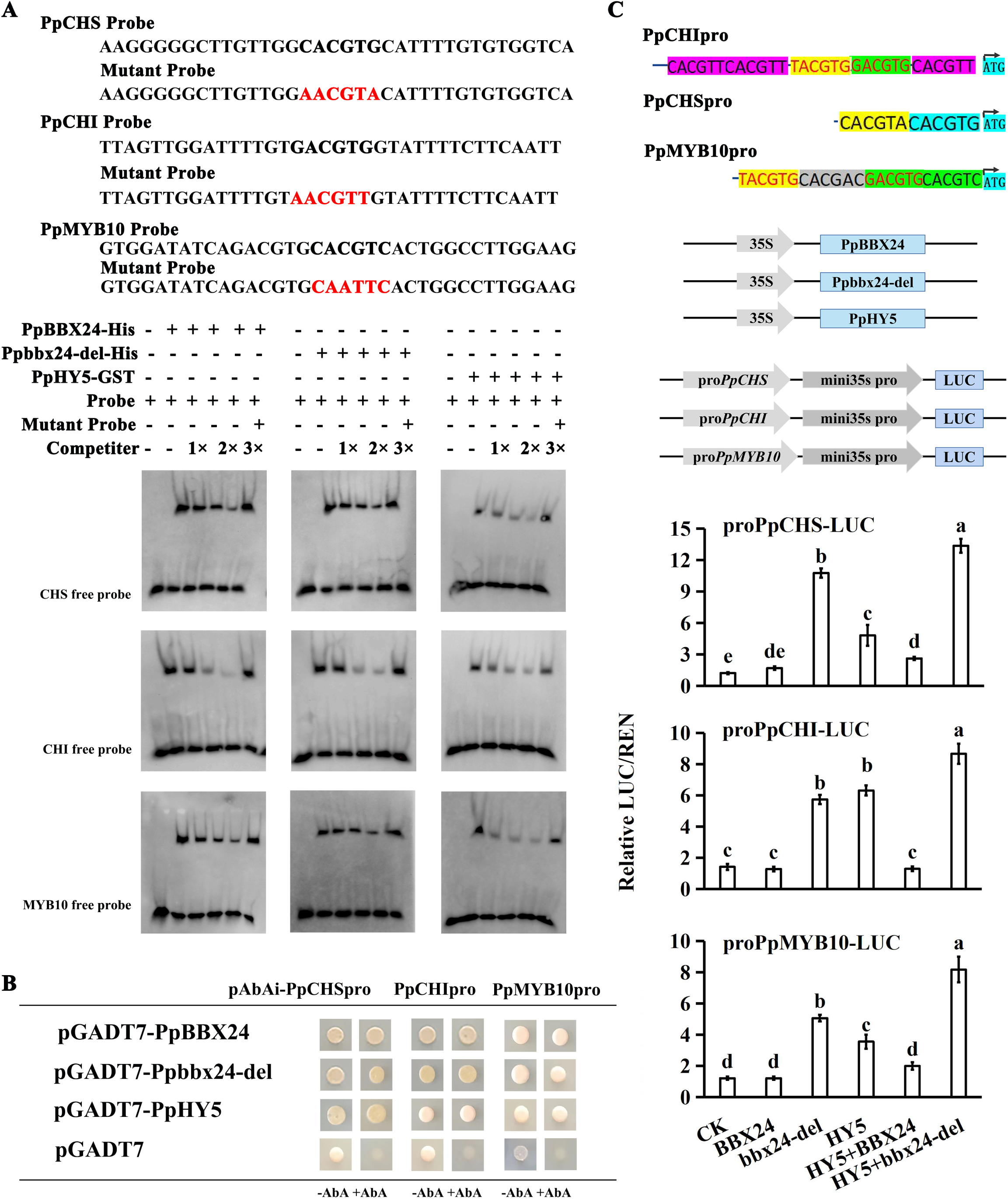
Binding of PpBBX24, Ppbbx24-del, and PpHY5 to G-box motifs in the *PpCHS*, *PpCHI*, and *PpMYB10* promoters. A, EMSA showing binding of PpBBX24, Ppbbx24-del, and PpHY5 to G-box motifs in *PpCHS*, *PpCHI*, and *PpMYB10* promoters. Probe: biotin-labeled fragment containing G-box motif, ‘competitor’ probe: non-labeled competitive probe. The mutant probe contained two nucleotide mutations. B, Yeast one-hybrid assays showing the interaction between PpBBX24, Ppbbx24-del, PpHY5, and promoters of *PpCHS*, *PpCHI*, and *PpMYB10*. C, Effects of PpBBX24, Ppbbx24-del, and PpHY5 on promoter activity of *PpCHS*, *PpCHI*, and *PpMYB10* as demonstrated by dual-luciferase assays.

To clarify the regulatory modes of PpBBX24, Ppbbx24-del, and PpHY5 on *PpCHS*, *PpCHI*, and *PpMYB10*, we fused their promoters to a LUC reporter gene for dual-luciferase assays. The observed high luciferase activity implied that Ppbbx24-del and PpHY5 could activate the *PpCHS*, *PpCHI*, and *PpMYB10* promoters. The activation effect of Ppbbx24-del on *PpCHS* and *PpMYB10* was significantly higher than that on PpHY5. Dual-luciferase assays showed that PpBBX24 had no significant influence on the promoter activities of *PpCHS*, *PpCHI*, and *PpMYB10*. Further analyses showed that Ppbbx24-del enhanced the effect of PpHY5 on *PpCHS*, *PpCHI*, and *PpMYB10* promoter activity, whereas PpBBX24 attenuated this effect. The luciferase activity was less than half as much as when PpHY5 was expressed alone, and the activation of *PpCHI* and *PpMYB10* genes was not significantly different from that of the control (Figure 7C). These results show that PpBBX24 inhibits the regulation of PpHY5 on structural genes and negatively regulates anthocyanin synthesis, whereas Ppbbx24-del strongly activates anthocyanin biosynthesis and does not depend on PpHY5.

## Discussion

### Ppbbx24-del is a potently positive regulator of anthocyanin biosynthesis in response to light

Anthocyanin accumulation is a complex process induced by environmental factors such as light and temperature, as well as internal factors during development (Xie et al., 2012; Fang et al., 2019b). Many BBX transcription factors, including AtBBX20, AtBBX21, AtBBX22 (Brusch et al., 2020), AtBBX23 (Zhang et al., 2017), AtBBX24 (Job et al., 2018) and AtBBX25 (Gangappa et al., 2013), have been found to regulate *HY5* transcription in *Arabidopsis*; in response to optical signals, they are associated with anthocyanin accumulation. The functions of MdBBX20 (Fang et al., 2019b), MdBBX22 (An et al., 2019), MdCOL4 (Fang et al., 2019a) and MdCOL11 (Bai et al., 2014), and other BBX transcription factors are affected by UV-B treatment in apples, as they regulate the expression of anthocyanin biosynthesis genes and ultimately affect anthocyanin levels. A total of 37 BBX family proteins have been identified in pears, among which PpBBX16, PpBBX18, and PpBBX21 are involved in anthocyanin synthesis (Bai et al., 2019a; Bai et al., 2019b; Talar et al., 2021). It was found that the expressions of *PpBBX16*, *PpBBX18* and *PpBBX21* in ‘Red Zaosu’ pear were significantly up-regulated then decreased and showed a fluctuating trend, after natural light treatment (Bai et al., 2019a; Bai et al., 2019b). Since the red variation of ‘Red Zaosu’ pear is mainly caused by the mutation of PpBBX24, is the expression of *PpBBX24* and *Ppbbx24-del* affected by light? This study detected the expression of *PpBBX24* and *Ppbbx24-del* in the skins of ‘Zaosu’ and ‘Red Zaosu’ at different times after bag-picking, it was found that their expressions tended to be up-regulated after bag-picking, but then fluctuated greatly, with no obvious regularity in the whole (Figure 1). These results indicated that the expressions of *PpBBX24* and *Ppbbx24-del* were not directly regulated by light. However, the fruit coloring of the ‘Red Zaosu’ pear was inhibited under the shading condition, indicating that light is the necessary condition for Ppbbx24-del to display its function.

### The deletion of the NLS and VP domains of Ppbbx24-del is the main reason for the changes **in its subcellular localization and PpHY5 interaction**

It is well known that transcription factors are required to achieve the regulation of gene expression in the nucleus (Hackenberg et al., 2012). In this study, the normal PpBBX24 protein was localized in the nucleus, consistent with the characteristics of normal transcription factors. However, the mutant Ppbbx24-del protein showed colocalization with the nucleoplasm with a weaker nuclear localization effect, which may be due to the absence of the nuclear localization sequence (NLS) domain in its C-terminus. After the C-terminus of AtBBX24 was truncated, an EGFP fusion protein (EGFP-N-BBX24) containing two B-box domains at the N-terminus of BBX24 was found in both the nucleus and cytoplasm (Yan et al., 2011). This was similar to the Ppbbx24-del mutation observed in this study (Figure 2). In the present study, Ppbbx24-del had a significant positive regulatory effect on the expression of *PpCHS*, *PpCHI*, and *PpMYB10*, further suggesting that it can enter the nucleus. Similar studies have been conducted previously. For example, the BZR1 transcription factor of *Arabidopsis* is localized in the cytoplasm; however, when the BR processing pathway is activated, BZR1 can be recruited into the nucleus after dephosphorylation (Wang et al., 2021). Some studies showed proteins that do not possess nuclear localization sequences can be assisted by some nuclear localization proteins to enter the nucleus. For example, FIT is located in the nucleus and can assist bHLH039, originally located in the cytoplasm, in entering the nucleus (Torfimov et al., 2019). OsPIE3 can modify the subcellular localization of PID2, thereby promoting its nuclear recruitment to the plasma membrane (Wang et al., 2023). The function of Ppbbx24-del in this study may also be achieved through some cofactors, but the specific reasons require further investigation.

Compared to PpBBX24, Ppbbx24-del lacks a valine-proline (VP) domain at the C-terminus. In *Arabidopsis*, the VP domain of CRY2 plays an important role in its interaction with HY5 (Ponnu et al., 2019). Previous studies have shown that BBX usually interacts with HY5 (An et al., 2019; Bursch et al., 2020), and we verified that PpBBX24 could interact with PpHY5, but Ppbbx24-del could not (Figure 5), which may be related to the deletion of the NLS and VP domains of Ppbbx24-del.

### PpBBX24 negatively regulated anthocyanin accumulation collaborated with PpHY5, while Ppbbx24-del positively regulated anthocyanin accumulation without PpHY5

Proteins of the same family may have different functions (Bursch et al., 2020). In *Arabidopsis*, phytochrome-interacting factors (PIFs) and LONG HYPOCOTYL IN FAR-RED1 (HFR1) belong to the 15th subgroup of the bHLH family, but are negative and positive regulatory factors of the light response, respectively (Shi et al., 2013; Xu et al., 2017). In the IV family of *Arabidopsis* BBX transcription factors, BBX20, BBX21, BBX22 (Bursch et al., 2020), and BBX23 (Zhang et al., 2017) were confirmed to be positive regulators of the light response, whereas BBX18 (Ding et al., 2018), BBX19 (Wang et al., 2015), BBX24 (Jiang et al., 2012) and BBX25 (Lyu et al., 2020a) were identified as negative regulatory factors. The C-terminal regions of different genes in the BBX family determine their functions. The gene functions of the positive regulator AtBBX21 and negative regulator AtBBX24 were reversed when their C-terminals were exchanged. The original negative regulator no longer inhibited downstream genes but promoted their expression (Job et al., 2018). The deletion of PpBBX24 resulted in a coding frame shift such that there was an early termination gene and the C-terminals were changed (Ou et al., 2020). Our study confirmed that Ppbbx24-del is associated with the red peel of pears, in contrast to PpBBX24.

HY5 is a basic leucine zipper (bZIP) transcription factor with powerful functions. Different photoreceptors mediate the light signal transduction by modulating HY5 activity and/or transcription to complete various life activities (Bhatia et al., 2021; Zhao et al., 2021b). In apples, MdMPK6 interacts directly with MdHY5 via phosphorylation under light conditions to regulate anthocyanin accumulation (Xing et al., 2023). MdWRKY11 binds to the promoter of *MdHY5* and promotes its expression (Liu et al., 2019b). In strawberries, FvbHLH9 interacts with FvHY5 to form HY5-bHLH heterodimers and activates the expression of the structural gene *FvDFR* (Li et al., 2020). HY5, together with BBXs, work in concert to control the expression of downstream target genes and regulate anthocyanin synthesis (An et al., 2019; Bursch et al., 2020). In *Arabidopsis*, AtBBX21 is a positive regulatory factor that combines with AtHY5 to regulate anthocyanin synthesis, whereas AtBBX24 is a negative regulatory factor that interferes with the binding of AtHY5 and the structural gene promoter of anthocyanin (Job et al., 2018). In apples, MdCOL4 interacts with MdHY5 to interfere with its binding to the anthocyanin structural gene promoter (Fang et al., 2019a). In this study, we confirmed that PpBBX24 interacts with PpHY5 both in vitro and in vivo, whereas the mutated PpbbX24-del did not interact with PpHY5 (Figure 5).

HY5 directly binds to the G-box motifs present in downstream target genes to activate their expression (Heng et al., 2019; Wang et al., 2020; Job et al., 2021; Li et al., 2022; Xing et al., 2023). These genes include *CHS*, *CHI*, *FLS*, *DFR*, and other structural genes related to anthocyanin synthesis (Li et al., 2020; Zhao et al., 2021b). HY5 can also indirectly influence anthocyanin synthesis by regulating the expression of MYB transcription factors and the MBW complex (Fang et al., 2019a). For example, in *Arabidopsis*, AtMYBD employed by AtHY5 increases anthocyanin accumulation by repressing *AtMYBL2* (Nguyen et al., 2015). AtHY5 can also regulate anthocyanin biosynthesis by binding to the *AtMYB75*/*AtPAP1* promoter and inducing transcriptional activation of transcription factors (Shin et al., 2013). In apples, MdHY5 binds to the G-box element in the *MdMYBDL1* promoter to activate its expression (Liu et al., 2019a). MdHY5 can also bind to E-box and G-box motifs to activate its own expression and *MdMYB10* (An et al., 2017). In this study, Ppbbx24-del did not interact with PpHY5 in vitro but enhanced the promoter activity of *PpCHS*, *PpCHI*, and *PpMYB10*. When Ppbbx24-del and PpHY5 coexisted, the activation effect accumulated and promoter activity was further enhanced. PpBBX24 significantly suppressed the promoter activities of *PpCHS*, *PpCHI*, and *PpMYB10* by forming a PpBBX24-PpHY5 dimer. This suggests that PpBBX24 and PpHY5 antagonize anthocyanin synthesis (Figure 7). Our results demonstrate that mutated PpBBX24 genes acquire the opposite function: PpBBX24 acts synergistically with PpHY5 to regulate the accumulation of pyocyanin as a negative regulatory factor, whereas Ppbbx24-del as a positive regulatory factor does not depend on PpHY5.

Red pear germplasm resources are rare; pears with red peel have received increasing attention from customers because of their bright appearance and sweet taste, and analysis of the mechanism of the red character is of great significance for red pear breeding (Zhai et al., 2021; Alabd et al., 2022). Based on the data presented herein, we summarized the working model of PpBBX24 and Ppbbx24-del in the regulation of anthocyanin accumulation under light conditions (Figure 8). Under normal light conditions, in normal cells, PpHY5 can bind to the G-box motifs of the *PpMYB10*, *PpCHS*, and *PpCHI* promoters and promote their transcription. PpBBX24 is located in the nucleus and can bind to the G-box in the promoters of the above genes but has no significant effect on the expression of these genes. The PpBBX24-PpHY5 protein interaction weakens the regulatory effects of these genes. PpBBX24 also competitively binds to the G-box motif of active PpHY5, which may inhibit the activation of active PpHY5 in downstream genes. In the mutant cells, approximately half of PpBBX24 was replaced by the mutant Ppbbx24-del. On the one hand, the more active PpHY5 was released. On the other hand, when Ppbbx24-del colocalized with the nucleoplasm; part of the Ppbbx24-del entering the nucleus can also regulate light-induced anthocyanin biosynthesis through direct activation of *PpMYB10*, *PpCHS,* and *PpCHI* promoters in red pear fruit. Therefore, the transcript levels of *PpMYB10*, *PpCHS*, and *PpCHI* were significantly higher in the mutant cells, which promoted anthocyanin synthesis and caused the cells to appear red. This study provides new insights into the regulation of light-induced anthocyanin biosynthesis in pear fruit at both the transcriptional and post-translational levels and provides an important theoretical reference for the molecular breeding of the red pear.

**Figure 8.**
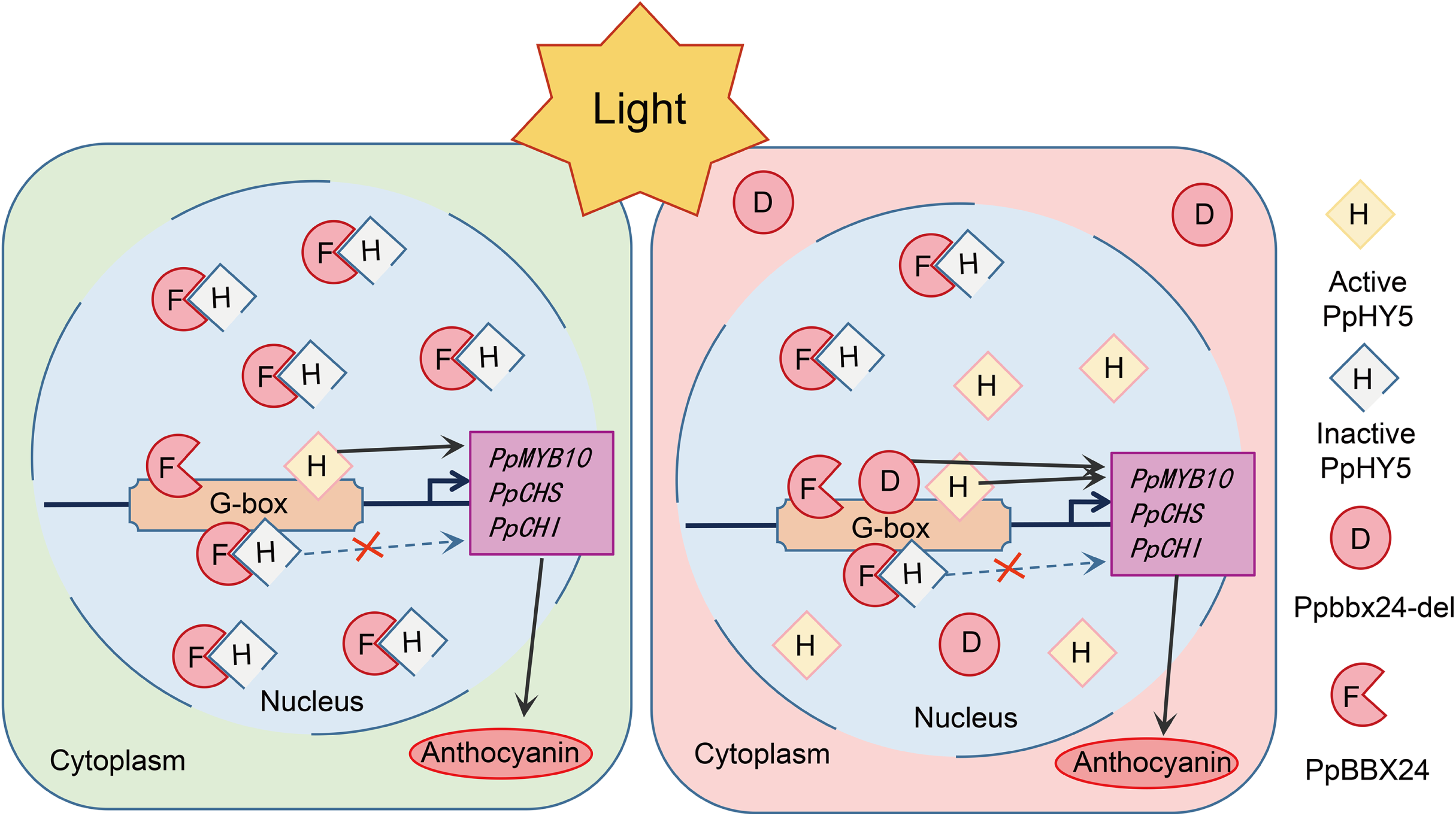
Prediction models of the biosynthesis of ‘Zaosu’ (left) and ‘Red Zaosu’ (right) by synergic regulation of PpBBX24 and Ppbbx24-del with PpHY5.

## Materials and Methods

### Plant materials

Young leaves and fruits were selected at random from ‘Zaosu,’ ‘Red Zaosu,’ and ‘Huang Guan’ trees. The young leaves of 30 red F1 plants and 30 green F1 plants of ‘Red Zaosu × Huang Guan’ (10 of which were mixed for biological replication) were collected for transcriptome and metabolome sequencing. Young fruits were bagged in double-layered paper bags 20 days after full blooming, and the bags were picked 21 days before fruit ripening. From the day of bag picking, five fruits were randomly picked every three days outside the crown until the fruits were ripe, and the peels were collected for anthocyanin content measurement and gene cloning. All the experiments were grown in the orchard of Research Institute of Pomology of Chinese Academy of Agricultural Sciences (Xingcheng, China, 120°44′38″E, 40°37′9″N), with routine fertilizer and water management.

The tobacco NC89 (*Nicotiana tabacum* L. var. NC89) vaccine for stable transformation was provided by our laboratory and was cultured conventionally on MS medium. The *Nicotiana benthamiana* cultured normally for approximately one month was used for transient expression.

### Measurement of anthocyanin content

The anthocyanins were extracted as described by Wan et al. (2017). Each peel sample (1 g) was extracted in 10 mL 1% HCl/CH_3_OH at 4 ℃ for 24 h in the dark. Absorbance at 553 nm and 600 nm were measured using a UV-vis spectrophotometer. OD_553 nm_–OD_600 nm_=0.01 was regarded as a unit to calculate the relative anthocyanin content.

### RNA isolation and real-time PCR

Total RNA was isolated using the EasySpin Plus Rapid Plant RNA Extraction Kit (RN38; Aidlab, Beijing, China), and first-strand cDNA was synthesized using PrimeScript RT Master Mix (Perfect Real Time) (RR036Q; TaKaRa, Ohtsu, Japan) following the manufacturer’s instructions. qPCR was performed using the Luna Universal qPCR Master Mix (M3003X; NEW ENGLAND Biolabs) on a Bio-Rad CFX96 Touch fluorescence quantitative PCR instrument. The RT-qPCR reaction was performed in a 20 μL system with the ACTIN gene (GU830958.1) as the internal reference, and the primers used are shown in Table S1. The cycle threshold (Ct) 2^-ΔΔCT^ method was used for the relative quantification of mRNA. The reaction was set up for three repetitions, and SPSS software was used to analyze the significance of the results.

### Subcellular localization analysis

Full-length coding sequences (CDSs), without termination codons, of PpBBX24 and Ppbbx24-del were independently cloned from the cDNA of the ‘Red Zaosu’ pear. They were connected to the vector pRI101, which contained the CaMV 35S promoter, and integrated with an EGFP tag (primers listed in Table S1). The empty vector p35SPPDK-EGFP-Flag was used as a control. *A. tumefaciens* GV3101 lines harboring the vectors were independently infiltrated into onion epidermis for 30 min. After dark culture on MS medium at 28 ℃ for two days, fluorescence was observed by a fluorescence microscope (IX51, with DP22 image acquisition system; Olympus, Japan).

### Transformation of tobacco with PpBBX24 and Ppbbx24-del

The full-length CDSs of *PpBBX24* and *Ppbbx24-del* were independently cloned into the pRI101 vector. *A. tumefaciens* EHA105 lines harboring the vectors (pRI101-*PpBBX24* and pRI101-*Ppbbx24-del*) were independently infiltrated into the leaves of NC89 tobacco for 10 min, and then cultured on medium (MS + TDZ 2.0 mg/L + NAA 0.5 mg/L, PH 5.8) for three days in the dark. Next, the leaves were transferred to MS medium containing 20 mgꞏL^-1^ kanamycin and cultured for two weeks in the dark until resistant buds regenerated. At least three positive buds were obtained for each gene and transplanted into the field after rooting. Target gene expression was detected by RT-PCR.

### Transient transformation analysis in pear fruit

A 418 bp fragment was selected from the Ppbbx24-del gene coding frame to design specific primers, which were inserted into both sides of the intron of the pKAN vector in the positive and negative directions, and then connected to the pART27 vector. *A. tumefaciens* GV3101 with pRI101-*PpBBX24*, pRI101-*Ppbbx24-del*, and RNAi-*Ppbbx24-del* vectors was injected into pear fruits. After infiltration, the fruits were kept under dark conditions for at least two days and light for three days before photos were taken. The fruit peels surrounding the injection sites were collected for gene expression and anthocyanin content detection.

### Transcriptome sequencing and metabolome analysis

Sample preparation, extraction, flavonoid identification and quantification, and RNA library preparation and sequencing were performed at Wuhan MetWare Biotechnology Co., Ltd. (www.metware.cn), using standard procedures previously described by Wang et al. (2019) and Tu et al. (2022). Sample analysis was performed according to Li et al. (2021), and the specific accumulation VIP value and fold change of metabolites were set at 1.0 and 2.0, respectively. Analysis of RNA sequencing data was carried out according to the method described by Tu et al. (2022), where the p-value and fold change were set at 0.05 and 2.0, respectively, after differential gene correction.

### Bimolecular fluorescence complementation assay (BiFC)

The CDSs, without the termination codon, of PpBBX24, Ppbbx24-del, and *PpHY5* were cloned into pSPYNE and pSPYCE vectors using a Seamless Cloning Kit (D7010S; Beyotime). Then *A. tumefaciens* GV3101 harboring the constructs was transiently co-expressed with all possible combinations of PpBBX24-NE/Ppbbx24-del-NE and *PpHY5-CE* fusion proteins in the onion epidermis. Fluorescence was observed using subcellular localization assays.

### Yeast two-hybrid assays

The Yeast Two-Hybrid assays were performed according to the Matchmaker Gold Yeast Two-Hybrid System (Cat#630489; Clontech) kit instructions. The coding sequences of *PpBBX24*, *Ppbbx24-del*, and *PpHY5* were connected to pGBKT7 or pGADT7 vectors. The vectors were then transformed into Y2HGold yeast cells using the polyethylene glycol/lithium acetate method. To screen for interactions, the cells were plated onto SD/-Trp/-Leu medium. The putative transformants were then transferred to SD/-Trp/-Leu/-His/-Ade/AbA/X-a-Gal medium. Colony growth and color changes were also observed.

### Pull-down assays

*PpBBX24* and *Ppbbx24-del* were cloned into a pET-N-His-TEV vector containing a His-tag sequence. PpHY5 was cloned into the pET-N-GST-PreScission vector using a GST tag sequence. The PpBBX24-His, Ppbbx24-del-His, and PpHY5-GST fusion proteins were individually expressed in and purified from *Escherichia coli* BL21 (DE3), respectively. The His-tag Protein Purification Kit (Denaturant-resistant) (P2229S; Beyotime) and GST-tag Protein Purification Kit (P2262; Beyotime) Ni columns were used for purification. Subsequently, PpBBX24-His and Ppbbx24-del-His were incubated with PpHY5-GST and GST, respectively, using a GST-tag Purification Resin. The presence of anti-His fusion proteins was detected using western blotting.

### Promoter cloning

Specific primers (Table S1) were designed according to the sequence information of pear reference genome ‘Zhongai-1’ pear (Ou et al., 2020). DNA of ‘Zaosu’ and ‘Red Zaosu’ were used as templates. About 2000 bp upstream of the start codon of *PpCHI* (Pdr1g010900), *PpCHS* (Pdr4g019560), *PpANS* (Pdr6g006790), *PpUFGT* (Pdr16g020840), *PpMYB10* (Pdr9g001390), and *PpHY5* (Pdr8g016720) genes were amplified, respectively. The sequence differences between the two varieties were compared. Plant CARE (http://bioinformatics.psb.ugent.be/webtools/plantcare/html/) online software was used to analyze each component of the promoter sequence.

### Electrophoretic mobility shift assays (EMSA)

Probes for each promoter containing G-box motifs were synthesized by Taihe Biotechnology Co., Ltd. The probe was labeled with biotin using an EMSA Probe Biotin Labeling Kit (GS008; Beyotime). The induced purification of prokaryotes was similar to that observed in pull-down assays. EMSA was performed according to the instructions of the Chemiluminescent EMSA Kit (GS009; Beyotime), and the results were imaged using a Bio-Macromolecule Imager (ChemiDoc; Bio-Rad, USA).

### Yeast one-hybrid assays

*PpBBX24*, *Ppbbx24-del*, and *PpHY5* were cloned into the pGADT7 vector, and the promoter fragments of *PpCHS*, *PpCHI*, and *PpMYB10* were cloned into the pAbAi vector. Y1HGold yeast cells were transfected with recombinant plasmids. Transformed cells were detected on SD/-Leu/AbA plates, and the binding ability of transcription factors to promoters was determined based on yeast growth.

### Dual-luciferase assays

The LUC reporter assays were conducted as described by Ye et al. (2017). The CDS of PpHY5 was recombined into the pRI101 vector, and the promoter fragments of *PpCHI*, *PpCHS*, and *PpMYB10* were inserted into the pGreenⅡ800-LUC vector. The *Agrobacterium* injection was used to express effector and reporter plasmids in tobacco leaves. The transfected leaves were incubated at 24 ℃ for four days, then the fluorescence activity was measured on enzyme calibration (SPECTRA MAX 190) using a Dual Luciferase Reporter Gene Assay Kit (RG027; Beyotime). The ratio of LUC to GUS activity was calculated to determine the promoter activity.

## Acknowledgments

We thank the anonymous reviewers for their comments on the manuscript.

## Figure legends

**Figure S1.** Coding sequence (CDS) alignment and translation of PpBBX24 and Ppbbx24-del in ‘Red Zaosu.’

